# PointSite: a point cloud segmentation tool for identification of protein ligand binding atoms

**DOI:** 10.1101/831131

**Authors:** Zhen Li, Xu Yan, Qing Wei, Xin Gao, Sheng Wang, Shuguang Cui

**Affiliations:** Chinese University of Hong Kong, Shenzhen (CUHKSZ); Shenzhen Research Institute of Big Data (SRIBD); King Abdullah University of Science and Technology (KAUST); Tencent AI Lab

**Keywords:** Protein ligand binding site (LBS), Ligand atoms, Point clouds segmentation, Submanifold Sparse Convolution

## Abstract

Accurate identifications of ligand binding sites (LBS) on protein structure is critical for understanding protein function and designing structure-based drug. As the previous pocket-centric methods are usually based on the investigation of pseudo surface points (PSPs) outside the protein structure, thus inherently cannot incorporate the local connectivity and global 3D geometrical information of the protein structure. In this paper, we propose a novel point clouds segmentation method, PointSite, for accurate identification of protein ligand binding atoms, which performs protein LBS identification at the atom-level in a protein-centric manner. Specifically, we first transfer the original 3D protein structure to point clouds and then conduct segmentation through Submanifold Sparse Convolution (SSC) based U-Net. With the fine-grained atom-level binding atoms representation and enhanced feature learning, PointSite can outperform previous methods in atom-IoU by a large margin. Furthermore, our segmented binding atoms can work as a filter on predictions achieved by previous pocket-centric approaches, which significantly decreases the false-positive of LBS candidates. Through cascaded filter and re-ranking aided by the segmented atoms, state-of-the-art performance can be achieved over various canonical benchmarks and CAMEO hard targets in terms of the commonly used DCA criteria. Our code is publicly available through https://github.com/PointSite.

## 1 Introduction

In computational biology, a fundamental question remains to be answered: given a protein structure, can we accurately identify the atoms that form the ligand-binding sites (LBS) [27]? In cellular environment, most proteins perform biological functions by interacting with other ligands, which are small molecules ranging from metal ions, organic or inorganic molecules, to polymers such as polysaccharides and short peptides [33]. The accurate identification of LBS on protein surfaces is critical for understanding the functions of the protein [19], and in turn an indispensable step for rational structure-based drug design (SBDD) [2]. However, it is highly expensive and time-consuming to detect the protein LBS using experimental techniques, which often requires solving the 3D structure of the protein-ligand complexes. Furthermore, although the number of 3D structures in protein data bank (PDB) grows rapidly, protein-ligand binding complex structure remain limited even if apo protein has solved structures [20]. Therefore, computational methods are valuable and needed to predict LBS on protein surface.

Formally, the protein LBS is defined as the heavy atoms on the protein that are within a certain distance (such as 6.5Å, denoted as definition radius) to any heavy atom of the ligand [6]. Till now, various approaches have been developed for the identification of protein LBS, which are comprehensively reviewed and summarized in [33, 48, 40, 32, 31, 15, 29, 9, 39, 4, 34, 46]. In general, the existing methods can be mainly categorized into two classes: template-based methods and template-free methods. In this work, we focus on template-free methods.

For template-free methods, we may divide them into pocket-centric and protein-centric ones (Fig. 1). The key principle of those pocket-centric approaches is based on the following two steps: (i) searching for pseudo surface points (PSPs) outside the protein structure, which could be regarded as the candidate LBS or pocket; (ii) identifying the binding atoms within a certain radius to these PSPs (denoted as identification radius). The existing pocket-centric methods can be categorized into geometric, energetic, and machine learning based on their main algorithmic strategy to search for PSPs [24]. The geometric strategy includes methods that identify the PSP by using the 3D geometric structure of the protein to search for cavities and pockets of the protein, which could be further classified into grid scanning (e.g., PocketPicker [42], LIGSITE [14], LIGSITE csc [19]), probe sphere (e.g., PASS [3]), alpha shape (e.g., CAST [30]), and alpha sphere (e.g., FPocket [28]); the energetic strategy includes methods that explore the energy of each position of the protein to the PSP (e.g., SiteHound [10], Q-sitefinder [10], PocketFinder [1]). Later, several machine learning methods have been proposed to exploit the protein features at residual and/or atomic level that influence the PSPs, and then use machine learning algorithms to identify (e.g, DeepSite [21] based on 3D-CNN) or rank (e.g., P2Rank [24] based on random forest) those PSPs.

**Fig. 1.**
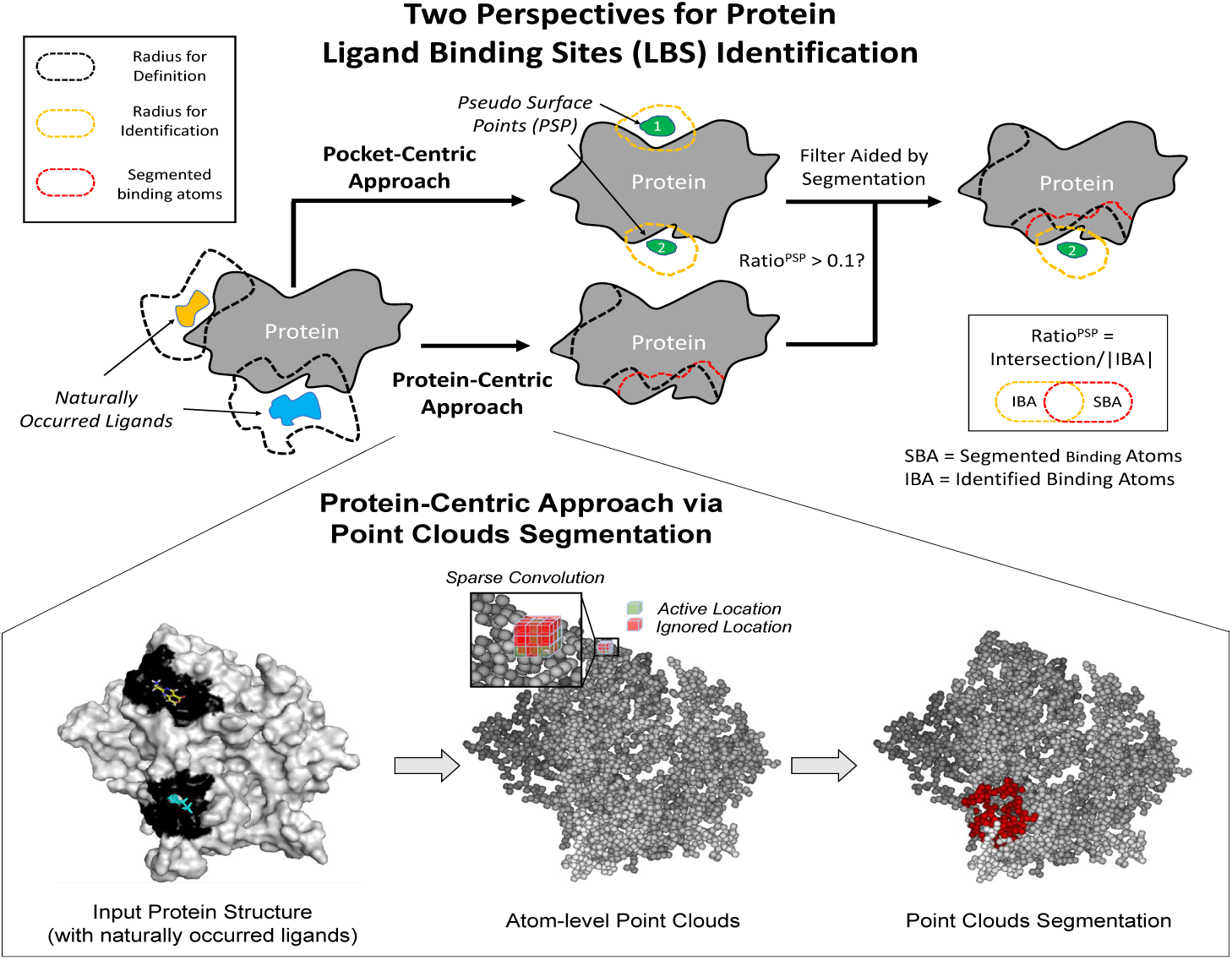
Given a protein structure, our purpose is to identify the binding atoms within a certain radius (i.e., definition radius) of the naturally occurred ligands. Two template-free perspectives for identifying protein ligand binding site (LBS) exist: pocket-centric and protein-centric. (i) The pocket-centric perspective firstly searches for pseudo surface points (PSPs) outside the protein structure, and then uses these PSPs to identify the protein binding atoms within a certain radius (i.e., identification radius); (ii) the protein-centric perspective directly identifies the binding atoms on the protein structure. The key idea of our approach is to formulate the binding atom identification problem as a point clouds segmentation problem, where each protein atom is represented as a point and the corresponding labels are non-binding or binding defined by the definition radius. Consequently, we show that our point clouds segmentation approach can serve as a powerful tool for prioritizing candidate PSPs identified by other pocket-centric approaches through filtering those PSPs that do not share enough common identified atoms with our approach. Best view in color.

It is clear that the pocket-centric methods answer the original question (i.e., identifying the ligand-binding atoms) in an indirect way. Furthermore, pocket-centric methods cannot explicitly take advantage of any connectivity between the atoms as well as the physicochemical characteristics of the atoms and/or residues on the protein structure. Therefore, a more direct way to identify LBS is the protein-centric perspective (Fig. 1). In this work, we present a point clouds segmentation approach for accurate identification of protein LBS at the atom-level. Firstly, we represent all the atoms in the query protein as point clouds, and mark those atoms in LBS as binding atom with label 1 and others with label 0. Then, we formulate the LBS identification problem as a point clouds segmentation problem, which could be elegantly solved by Sub-manifold Sparse Convolutional networks (SSCN) [11]. Considering conventional 3D convolutional neural network (3D-CNN) requires a large amount of memory consumption for dealing with 3D point clouds, sliding windows and down-sampling operations are exploited in DeepSite [21]. Such operations cannot take all point clouds as inputs and inherently harm the context learning. More seriously, traditional 3D-CNN suffers the submanifold dilation issue [11]. On the contrary, SSCN will keep the sparsity of the input points with each convolutional layer, and mark them to be either active or ignored (the lower panel in Fig. 1). Thus, such a deep neural network framework can input the entire point clouds at once and explicitly take into account the connectivity between the atoms as well as the physicochemical features of the protein atoms and residues simultaneously. Furthermore, SSC based U-Net learns the global and geometrical information of the query protein via the incrementally increased receptive field and skip connections, which in turn can capture the very complex relationship between the 3D protein structure and the binding atoms. Finally, our deep learning approach is a data-driven method, which can efficiently learn the characteristics of the entire database. Once the PointSite model has been trained, the inference time of our method is extremely fast (less than 1s per query protein) and the entire package is light-weighted.

Comprehensive experiments show that the proposed method, PointSite, achieved the state-of-the-art performance under the atom-level IoU (Intersection over Union) measurement. Moreover, our approach could serve as a tool for prioritizing candidate PSPs identified by other pocket-centric approaches through filtering those PSPs that do not share common identified atoms with our segmentation (described in Fig. 1). In terms of DCA (distances between the center of the Top-N identified LBS and any heavy atom of the naturally occurred ligand), the *de facto* gold standard pocket-centric measurement [24, 6], PointSite can significantly outperform all pocket-centric approaches by a large margin. For hard targets, such as the proteins with novel fold from CAMEO [12], the steadily better performance of PointSite not only confirms the generalization ability of our point clouds segmentation approach, but also proves the fact that the combination of the two complemental perspectives, protein-centric and pocket-centric approach, can lead to even better LBS identification.

## 2 Methods

In this paper, we propose a novel point clouds segmentation method, PointSite, for accurate identification of protein ligand binding atoms, which conducts protein LBS identification at atom-level in a protein-centric manner. Specifically, our proposed model mainly consists of three modules: 1) the atom point clouds transformation (APCT) module; 2) the ligand-binding atoms prediction (LAP) module; and 3) the LBS identification (LBSI) module. In the APCT module, we first transfer the original protein structure from the PDB format to atom-level point clouds. Taking into the transferred atom point clouds with simple features, the LAP module conducts ligand atoms prediction by utilizing a Submanifold Sparse Convolution (SSC) based U-net [35, 7]. Thanks to the sparse convolution operations, LAP can not only take all point clouds into consideration, but also contributes to a better global context learning. With the assistance of segmented ligand atoms, the accurate LBS identification is accomplished by filtering and re-ranking the prediction results generated by pocket-centric approaches.

### 2.1 Definitions and Problem Formulation

Given the PDB file of a query protein, our goal is to identify all the possible ligand-binding atoms, as well as output all possible LBS in a ranked list with the geometric center of each LBS. Note that our method is a template-free approach which only takes the original PDB file as the input. To identify LBS, our methods requires an additional pocket-centric approach to generate candidate LBS in a ranked order.

### 2.2 Atom Point Clouds Transformation (APCT) Module

Considering that previous pocket-centric approaches will predefine ad-hoc pseudo surface points (PSPs) for candidate LBS, here we conduct protein LBS identification at atom-level in a protein-centric perspective. Thus, we first transfer the original protein to atom point clouds. Given proteins with known ligand dataset, e.g., scPDB [23], all information has been presented in the original PDB file, including protein atoms and coordinates, ligand atoms and coordinates, protein residues and so on. In this paper, we transfer the original PDB file to the atom point clouds through a self-developed software LIG Tool (please refer to Supplemental Materials for detail). By taking a PDB file as input, the output is the features for each atom from the protein itself, including 21 residue type (A,R,N,D,C,Q,E,G,H,I,L,K,M,F,P,S,T,W,Y,V,UNK), 5 atom type (C,N,O,S,UNK) and the 3 coordinates for each atom. On the other hand, we label a protein atom as the ligand-binding atom (label 1) if it is within a sphere with the definition radius *r*_*def*_ (i.e., 6.5Å) to any heavy atom of the ligands, otherwise the label is 0. Under this setting, the features for all atoms is a 29-dimension vector, and the ground-truth label for each atom is 0/1.

In literature, the definition radius *r*_*def*_ ranges from 4.0Å to 6.5Å, and here we use 6.5Å for the following reasons: (a) this definition is used in scPDB that is an annotated database of druggable binding sites from PDB [23]; (b) the 6.5Å radius will consider those atoms that are in contact with binding atoms in implicit interaction with the ligands [37]; and (c) using this radius to define the ground-truth binding atoms will alleviate the class-imbalance issue.

### 2.3 Ligand Atoms Prediction (LAP) Module

In the APCT module, the input features and ground-truth labels of each protein atom have been defined. Considering that the atom number for each protein is usually over 10k, the submainfold dilation would occur if exploiting conventional 3D-CNN on voxelization representation. This will substantially reduce the sparsity of the input points from the previous convolutional layer and “blur” the output in the next layer (check an example in Fig. 2 in [11]). More seriously, conventional 3D-CNN cannot take entire point clouds as the input due to memory limitation. Such phenomenon is extremely disadvantageous for the context learning and identification of LBS. Inspired by submainfold sparse convolution (SSC) and the sparsity property of protein atoms, we exploit SSC based U-Net (Fig. 2) for ligand-binding atom prediction. Such architecture can not only take all point clouds as input for effectiveness, but also capture better global context and the geometric information of the protein structure.

**Fig. 2.**
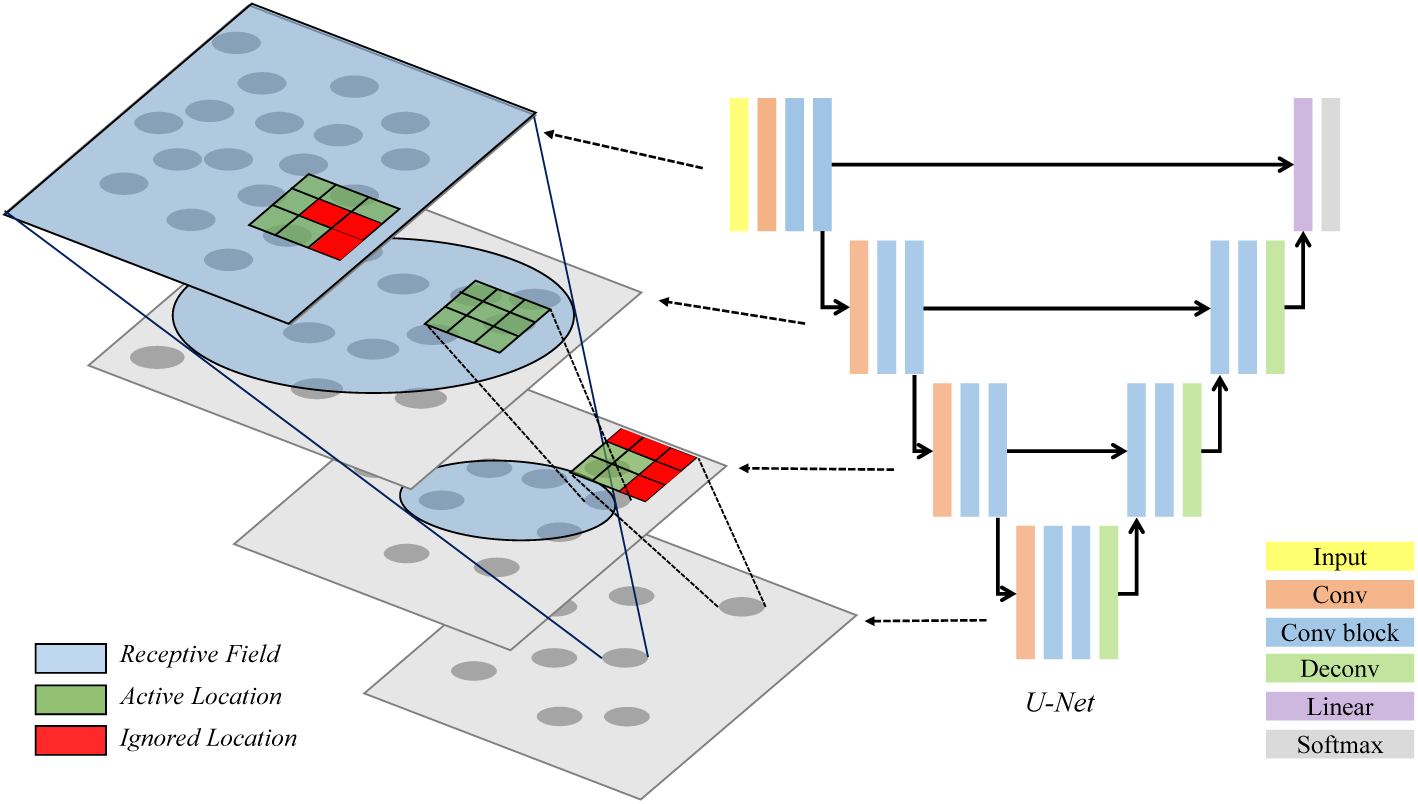
Submanifold Sparse Convolution based U-Net to avoid submanifold dilation.

### 2.4 LBS Identification (LBSI) Module

As mentioned above, pocket-centric approaches usually require additional information (e.g., geometric and/or energetic patterns of the ligand-binding pocket outside the protein) to define the PSP for the downstream LBS identification, while our point clouds segmentation method leverages native information of the atoms in the protein structure. Thus, it inspires us to design a merged model for LBSI through combining pocket-centric and protein-centric perspectives.

#### Filter aided by segmented binding atoms

The output of a certain pocket-centric approach (e.g., FPocket, and SiteBound) is a ranked list of candidate PSP, each of which consists of a cluster of PSPs that satisfy the corresponding criteria (such as high-scoring alpha spheres [28] and the surface points with high interaction energy [16]). Then starting from the Top-K candidate PSPs, we identify the ligand-binding atoms for a certain PSP with a certain identification radius *r*_*iden*_ in the same procedure as defining the ground-truth ligand-binding atoms. Therefore, for the *i*-th candidate PSP, the following filter strategy is used with the aid of our segmented ligand-binding atoms:

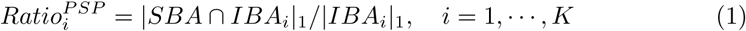

where SBA stands for segmented binding atoms, IBA_*i*_ stands for identified binding atoms for the *i*-th candidate PSP and *|·|*_1_ stands for the L1 norm (i.e., atom numbers). If the 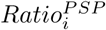 is below a threshold, then the *i*-th candidate PSP will be filtered out. Consequently the Top-K identifications would reduce to Top-K_1_ identifications for the pocket-centric approach.

#### Re-ranking by common binding atoms

From the above filtered Top-K_1_ identifications, we may further conduct a very simple re-ranking strategy based on the common binding atoms. Specifically, based on our SBA, we re-rank the remaining K_1_ identifications according to the common atoms as follows:

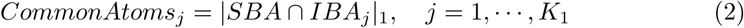

where SBA stands for segmented binding atoms set, IBA_*j*_ stands for identified binding atoms from the *j*-th filtered candidate PSP. Such simple strategy can not only benefit the protein-centric criteria (e.g, DCA), but also improves the performance of atom-level IoU.

## 3 Experiment

### 3.1 Datasets

There are three types of datasets used in this work: training, testing, and blind test, as shown in Tab. 1. We train and validate our method on the training dataset. The test dataset is applied for testing our method and comparing it with other approaches. We use CAMEO as the blind test dataset to show the performance on real-world cases. Each dataset shall belong to either chain-level or protein-level, where the former indicates that the entries in the dataset are protein chains while the latter are the protein complexes consisting multiple chains. It should be noted that for all datasets, we remove those entries with more than 5 binding ligands as well as those invalid ligands defined according to MOAD [41]. Supplemental Material gives more details on how we define the binding ligand using our in-house method LIG_Tool.

#### Training dataset

The dataset used to train our proposed method contains 6151 entries at chain-level, which is a subset of scPDB [8] create in 2017. Note that these entries share no more than 25% sequence identity with any test dataset.

#### Testing dataset

To evaluate our approach and compare with other approaches, we collect a set of well-recognized LBS datasets from available literatures [24]. These datasets include B277 [19], DT198 [47], ASTEX85 [13], CHEN251 [6], COACH420 [44, 36], and a large dataset HOLO4k [38]. See Supplemental Material for more details.

#### Blind test

To show the performance of our approach on “real-world” hard cases, we extracted a subset of the CAMEO [12] from 9/22/2018 to 9/22/2019 that contains 103 entries at chain-level, which has limited structural similarity (defined as *TMscore <* 0.6 [43]) and no sequence identity (*<* 25%) to our training data. After LIG Tool selection, we have 81 valid entries at chain-level in this dataset.

### 3.2 Comparison Methods and Evaluation Criteria

#### Comparison Methods

We compare out method PointSite with the following template-free pocket-centric methods: FPocket, SiteHound, MetaPocket2, DeepSite, and P2Rank on the test dataset. FPocket (version 3) is a typical geometric method based on filtering and clustering of alpha spheres found by Voronoi tessellation [28]. It is very popular and *de facto* the best geometric approach [24]. SiteHound is a typical energetic method based on the interaction energies between the protein and a PSP [16], and could also be regarded as the best energetic approach [24]. MetaPocket2 [18, 47] [47, 18] is a meta server to identify LBS based on the identification results from the eight popular pocket-centric methods: LIGSITEcsc [19], PASS [3], Q-SiteFinder [26], SURFNET [25], Fpocket (version 1) [28], GHECOM [22], ConCavity [5] and POCASA [45]. DeepSite [21] and P2Rank [24] are two recently developed approaches based on machine learning, where the former is rooted in 3D-CNN and the latter employs random forest. It should be noted that FPocket and P2Rank are open-source software, SiteHound has the stand-alone version, while MetaPocket2 and DeepSite only have the server version.

### 3.3 Evaluation Criteria

We use atom-level intersection over union (atom-IoU) to measure the performance on the accuracy of binding atoms identification. In particular, atom-IoU is defined as the intersection over union of two point sets, where the former point set is the identified ligand binding atoms and the latter one is the ground-truth ligand binding atoms. It is obvious that such metric is a typical protein-centric criterion, and all comparing methods are pocket-centric criteria, which require an identification radius (see Fig. 1) to identify the binding atoms from the PSPs. As the size, shape, and number of PSPs identified by different pocket-centric approaches are quite different, here we only select Top-N PSPs where N is the number of naturally occurred ligands, and output the maximal value of atom-IoU generated by 4.5Å, 5.5Å, and 6.5Å identification radius. We use DCA to show the original performance of all the pocket-centric approaches, as well as the performance in consideration of our segmentation results. Specifically, DCA is defined as the minimal distance between the geometric center of the Top-N identified LBS and any heavy atom of the naturally occurred ligands [6]. Again, N is the number of naturally occurred ligands. Usually, an identified LBS is considered as correct if the DCA is no further than 4Å.

### 3.4 Performance

#### Protein-centric Performance

We have extensively evaluated the protein-centric performance in terms of atom-IoU of PointSite and compared it against several widely used pocket-centric approaches on a variety of test datasets. As shown in Tab. 2, on almost all datasets, our point clouds segmentation method significantly outperforms all pocket-centric approaches including geometric methods, energetic ones, or machine learning or deep learning-based ones. P2Rank is the second best method. The only exception is the dataset CHEN251 which is the training set for P2Rank. However, even on this dataset, our result of 0.54 is comparable to that of P2Rank, which is 0.56. For all other datasets, our approach is 7%, 11%, 14%, 14%, and 15% better than P2Rank on DT198, B277, ASTEX85, COACH420, and HOLO4k, respectively. Note that P2Rank is the current state-of-the-art pocket-centric approach for LBS indentation, which is about 10% better than all the previous methods. These results indicate that our approach can significantly improve the accuracy of the binding atoms identification.

#### Pocket-centric Performance

As our method PoinstSite is a protein-centric approach, it is not easy to directly compare with the other pocket-centric approach in terms of DCA due to the difficulty in clustering the atom-level segmentation into several ranked classes in order to calculate DCA. However, the significantly improved identification results in terms of atom-IoU indicate that our approach could serve as a tool for prioritizing candidate PSPs (or, LBS) identified by other pocket-centric approaches through filtering those PSPs that do not share enough common identified atoms with our segmentation. Tab. 3 confirms this claim that our approach could significantly enhance the identification success rate measured by DCA for all pocket-centric approaches to a great deal on almost all the datasets (expect CHEN251 which is the training set for P2Rank). One notable example is the merged result of SiteHound. The original result of SiteHound is about 10-20% worse than that of P2Rank. However, the merged results of SiteHound is 4%, 2%, 11%, 8%, and 11% better than the original result of P2Rank on B277, DT198, ASTEX85, COACH420, and HOLO4k, respectively. In some datasets, such as B277 (1%), ASTEX85 (6%), COACH420 (5%), HOLO4k (6%), the merged results of SiteHound is even better than the merged result of P2Rank (value is shown in parenthesis), respectively. Such trends also appear in the measurement of atom-IoU (see Supplemental Tab. 2 to 6), and in some datasets the results are even better than that of PointSite itself. Therefore, the atom-level segmentation of PointSite can help the ranking and filtering of the identified LBS of the pocket-centric approaches, which significantly improve the performance not only in pocket-centric metric DCA but also protein-centric metric atom-IoU.

#### Blind Test

A natural question to ask is whether our point clouds segmentation approach learns the complex relationship between the 3D protein structure and the binding atoms or just simply “remembers” the training data. Although we removed all redundant and homologous proteins in the training data scPDB in the six testing datasets, it is still not sufficient to completely address this concern. To this end, we challenged our method on “real-world” hard targets from CAMEO that not only have no sequence identity (*<* 25%) but also have limited structural similarity (*TMscore <* 0.6 [43]) to our training data. Furthermore, these targets were released after 9/22/2018, which is even later than the latest version of scPDB released in 2017.

As shown in Table 4, among these 81 hard targets, our point clouds segmentation tool reaches 0.43 atom-IoU, which is 18%, 23%, 19%, 21%, and 6% better than FPocket, SiteHound, MetaPocket2, DeepSite, and P2Rank, respectively. When merged with our segmentation, the identification accuracy of all pocket-centric approaches is significantly improved in terms of both atom-IoU and DCA. An interesting discovery is that such improvement occurred more often in FPocket and SiteHound, which is a typical geometric and energetic approach, but not those consensus approach (say, MetaPocket2) or machine learning approaches (say, DeepStie and P2Rank). These results further prove the generalization power of the combination of our point clouds segmentation in protein-centric perspective with those individual unsupervised learning single approaches in pocket-centric perspective, because the two perspectives contain complement information to each other.

**Table 1.** Datasets used in this work.

|  | *scPDB* | *B277* | *DT198* | *ASTEX85* | *CHEN251* | *COACH420* | *HOLO4k* | *CAMEO* |
| --- | --- | --- | --- | --- | --- | --- | --- | --- |
| Purpose | Train | Test | Test | Test | Test | Test | Test | Blind |
| Type | Chain | Protein | Chain | Protein | Chain | Chain | Protein | Chain |
| Original Size | 16612 | 277 | 198 | 85 | 251 | 420 | 4543 | 103 |
| Actual Size | 6151 | 263 | 163 | 82 | 238 | 402 | 4063 | 81 |

**Table 2.** Comparison of identification performance on B277, DT198, ASTEX85, CHEN251, COACH420 and HOLO4K datasets in terms of atom-IoU (%).

| Method | *B277* | *DT198* | *ASTEX85* | *CHEN251* | *COACH420* | *HOLO4K* |
| --- | --- | --- | --- | --- | --- | --- |
| FPocket | 31.5 | 23.2 | 34.1 | 25.4 | 30.0 | 30.5 |
| SiteHound | 36.4 | 23.1 | 38.9 | 29.4 | 34.9 | 34.5 |
| MetaPocket2 | 37.3 | 25.8 | 37.5 | 32.8 | 37.7 | 38.8 |
| DeepSite | 34.0 | 29.1 | 37.4 | 27.4 | 33.9 | 33.2 |
| P2Rank | 49.8 | 38.6 | 47.4 | **56.5** | 45.3 | 48.8 |
| PointSite | **60.9** | **45.4** | **61.3** | 54.2 | **59.6** | **63.4** |

**Table 3.**
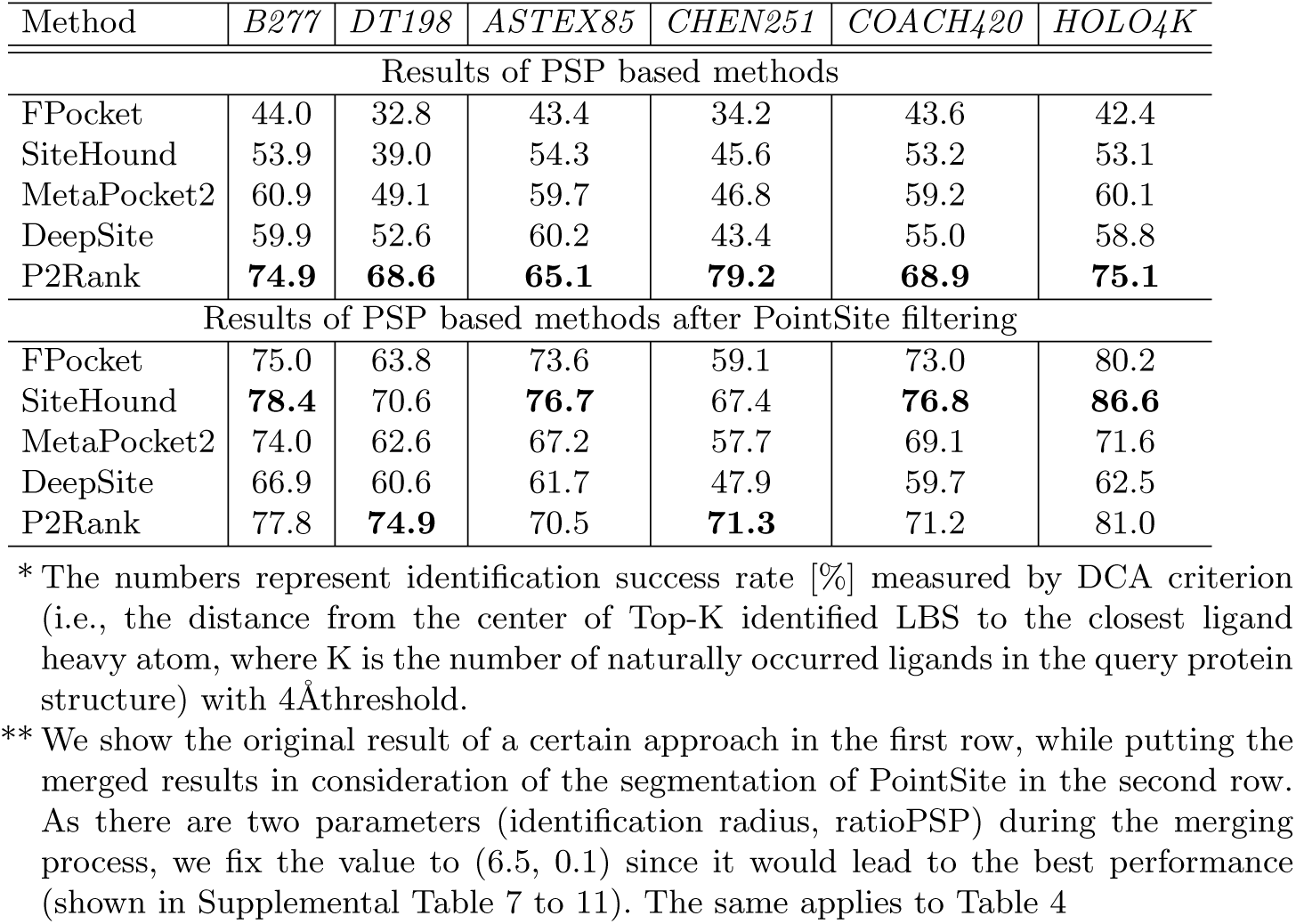
Comparison of identification performance on B277, DT198, ASTEX85, CHEN251, COACH420 and HOLO4K datasets in terms of DCA using the original results of those pocket-centric approaches as well as the merged results in consideration of the segmentation of PointSite.

| Method | *B277* | *DT198* | *ASTEX85* | *CHEN251* | *COACH420* | *HOLO4K* |
| --- | --- | --- | --- | --- | --- | --- |
| Results of PSP based methods |  |  |  |  |  |  |
| FPocket | 44.0 | 32.8 | 43.4 | 34.2 | 43.6 | 42.4 |
| SiteHound | 53.9 | 39.0 | 54.3 | 45.6 | 53.2 | 53.1 |
| MetaPocket2 | 60.9 | 49.1 | 59.7 | 46.8 | 59.2 | 60.1 |
| DeepSite | 59.9 | 52.6 | 60.2 | 43.4 | 55.0 | 58.8 |
| P2Rank | **74.9** | **68.6** | **65.1** | **79.2** | **68.9** | **75.1** |
| Results of PSP based methods after PointSite filtering |  |  |  |  |  |  |
| FPocket | 75.0 | 63.8 | 73.6 | 59.1 | 73.0 | 80.2 |
| SiteHound | **78.4** | 70.6 | **76.7** | 67.4 | **76.8** | **86.6** |
| MetaPocket2 | 74.0 | 62.6 | 67.2 | 57.7 | 69.1 | 71.6 |
| DeepSite | 66.9 | 60.6 | 61.7 | 47.9 | 59.7 | 62.5 |
| P2Rank | 77.8 | **74.9** | 70.5 | **71.3** | 71.2 | 81.0 |

\* The numbers represent identification success rate [%] measured by DCA criterion (i.e., the distance from the center of Top-K identified LBS to the closest ligand heavy atom, where K is the number of naturally occurred ligands in the query protein structure) with 4Å threshold.
\*\* We show the original result of a certain approach in the first row, while putting the merged results in consideration of the segmentation of PointSite in the second row. As there are two parameters (identification radius, ratioPSP) during the merging process, we fix the value to (6.5, 0.1) since it would lead to the best performance (shown in Supplemental Table 7 to 11). The same applies to Table 4

**Table 4.**
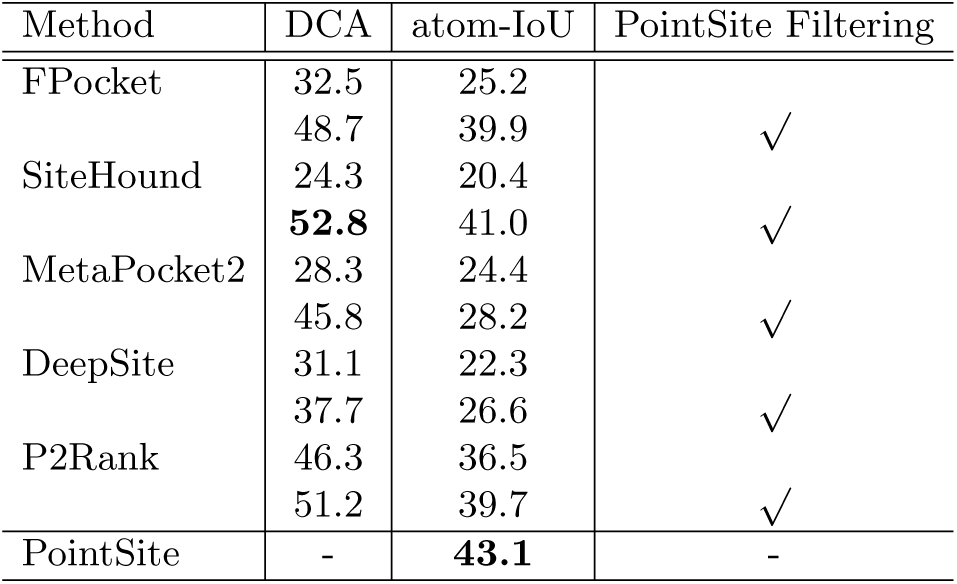
Comparison of identification performance on CAMEO blind datasets in terms of DCA criterion at 4Åthreshold as well as atom-IoU (%).

## 4 Conclusion and Discussion

We introduced PointSite, a novel protein-centric approach for accurate identification of ligand binding atoms, which significantly improves the atom-level IoU over all previous approaches by a great margin. Our method distinguishes itself from previous protein-centric approaches in that we formulate the binding atom identification problem as a typical point clouds segmentation problem. Nonetheless, all current available protein-centric methods are sequence-based approaches, which typically focus on the residue sequence of a protein to identify LBS. Therefore, these sequence-based methods are not able to explicitly capture the connectivity between the atoms on the proteins structure, but also unable to consider the 3D geometric property of the query protein. Moreover, they can only identify the binding residues instead of binding atoms. That’s why if the protein structure is available, these sequence-based approaches can only provide limited information compared to those pocket-centric methods [17], which is also the reason why we did not compare with sequence-based methods in this work

We have shown that the segmented binding atoms from PointSite could serve as a post processing tool to guide any pocket-centric approaches through a filtering and re-ranking strategy to prioritize the identified candidate PSPs. Since pocket-centric approaches might output many false positive results, a subsequent prioritization step can greatly boost the performance of such tools. This is proved by the evidence that the merged results can greatly enhance the identification accuracy in terms of commonly used DCA criterion as well as the atom-level IoU for all pocket-centric approaches.

## Supporting information

Supplemental Materials for PointSite

## References

1. An, J., Totrov, M., Abagyan, R.: Pocketome via comprehensive identification and classification of ligand binding envelopes. Molecular & Cellular Proteomics 4(6), 752–761 (2005)

2. Anderson, A.C.: The process of structure-based drug design. Chemistry & biology 10(9), 787–797 (2003)

3. Brady, G.P., Stouten, P.F.: Fast prediction and visualization of protein binding pockets with pass. Journal of computer-aided molecular design 14(4), 383–401 (2000)

4. Broomhead, N.K., Soliman, M.E.: Can we rely on computational predictions to correctly identify ligand binding sites on novel protein drug targets? assessment of binding site prediction methods and a protocol for validation of predicted binding sites. Cell biochemistry and biophysics 75(1), 15–23 (2017)

5. Capra, J.A., Laskowski, R.A., Thornton, J.M., Singh, M., Funkhouser, T.A.: Predicting protein ligand binding sites by combining evolutionary sequence conservation and 3d structure. PLoS computational biology 5(12), e1000585 (2009)

6. Chen, K., Mizianty, M.J., Gao, J., Kurgan, L.: A critical comparative assessment of predictions of protein-binding sites for biologically relevant organic compounds. Structure 19(5), 613–621 (2011)

7. Çiçek, Ö., Abdulkadir, A., Lienkamp, S.S., Brox, T., Ronneberger, O.: 3d u-net: learning dense volumetric segmentation from sparse annotation. In: International conference on medical image computing and computer-assisted intervention. pp. 424–432. Springer (2016)

8. Desaphy, J., Bret, G., Rognan, D., Kellenberger, E.: sc-pdb: a 3d-database of ligandable binding sites—10 years on. Nucleic acids research 43(D1), D399–D404 (2014)

9. Fauman, E.B., Rai, B.K., Huang, E.S.: Structure-based druggability assessment—identifying suitable targets for small molecule therapeutics. Current opinion in chemical biology 15(4), 463–468 (2011)

10. Ghersi, D., Sanchez, R.: Easymifs and sitehound: a toolkit for the identification of ligand-binding sites in protein structures. Bioinformatics 25(23), 3185–3186 (2009)

11. Graham, B., Engelcke, M., van der Maaten, L.: 3d semantic segmentation with submanifold sparse convolutional networks. In: Proceedings of the IEEE Conference on Computer Vision and Pattern Recognition. pp. 9224–9232 (2018)

12. Haas, J., Barbato, A., Behringer, D., Studer, G., Roth, S., Bertoni, M., Mostaguir, K., Gumienny, R., Schwede, T.: Continuous automated model evaluation (cameo) complementing the critical assessment of structure prediction in casp12. Proteins: Structure, Function, and Bioinformatics 86, 387–398 (2018)

13. Hartshorn, M.J., Verdonk, M.L., Chessari, G., Brewerton, S.C., Mooij, W.T., Mortenson, P.N., Murray, C.W.: Diverse, high-quality test set for the validation of protein-ligand docking performance. Journal of medicinal chemistry 50(4), 726–741 (2007)

14. Hendlich, M., Rippmann, F., Barnickel, G.: Ligsite: automatic and efficient detection of potential small molecule-binding sites in proteins. Journal of Molecular Graphics and Modelling 15(6), 359–363 (1997)

15. Henrich, S., Salo-Ahen, O.M., Huang, B., Rippmann, F.F., Cruciani, G., Wade, R.C.: Computational approaches to identifying and characterizing protein binding sites for ligand design. Journal of Molecular Recognition: An Interdisciplinary Journal 23(2), 209–219 (2010)

16. Hernandez, M., Ghersi, D., Sanchez, R.: Sitehound-web: a server for ligand binding site identification in protein structures. Nucleic acids research 37(Suppl 2), W413–W416 (2009)

17. Hu, X., Wang, K., Dong, Q.: Protein ligand-specific binding residue predictions by an ensemble classifier. BMC bioinformatics 17(1), 470 (2016)

18. Huang, B.: Metapocket: a meta approach to improve protein ligand binding site prediction. OMICS A Journal of Integrative Biology 13(4), 325–330 (2009)

19. Huang, B., Schroeder, M.: Ligsite csc: predicting ligand binding sites using the connolly surface and degree of conservation. BMC structural biology 6(1), 19 (2006)

20. Jiang, M., Li, Z., Bian, Y., Wei, Z.: A novel protein descriptor for the prediction of drug binding sites. BMC bioinformatics 20(1), 1–13 (2019)

21. Jiménez, J., Doerr, S., Martínez-Rosell, G., Rose, A.S., De Fabritiis, G.: Deepsite: protein-binding site predictor using 3d-convolutional neural networks. Bioinformatics 33(19), 3036–3042 (2017)

22. Kawabata, T.: Detection of multiscale pockets on protein surfaces using mathematical morphology. Proteins: Structure, Function, and Bioinformatics 78(5), 1195–1211 (2010)

23. Kellenberger, E., Muller, P., Schalon, C., Bret, G., Foata, N., Rognan, D.: sc-pdb: an annotated database of druggable binding sites from the protein data bank. Journal of chemical information and modeling 46(2), 717–727 (2006)

24. Krivák, R., Hoksza, D.: P2rank: machine learning based tool for rapid and accurate prediction of ligand binding sites from protein structure. Journal of cheminformatics 10(1), 39 (2018)

25. Laskowski, R.A.: Surfnet: a program for visualizing molecular surfaces, cavities, and intermolecular interactions. Journal of molecular graphics 13(5), 323–330 (1995)

26. Laurie, A.T., Jackson, R.M.: Q-sitefinder: an energy-based method for the prediction of protein–ligand binding sites. Bioinformatics 21(9), 1908–1916 (2005)

27. Laurie, R., Alasdair, T., Jackson, R.M.: Methods for the prediction of protein-ligand binding sites for structure-based drug design and virtual ligand screening. Current Protein and Peptide Science 7(5), 395–406 (2006)

28. Le Guilloux, V., Schmidtke, P., Tuffery, P.: Fpocket: an open source platform for ligand pocket detection. BMC bioinformatics 10(1), 168 (2009)

29. Leis, S., Schneider, S., Zacharias, M.: In silico prediction of binding sites on proteins. Current medicinal chemistry 17(15), 1550–1562 (2010)

30. Liang, J., Woodward, C., Edelsbrunner, H.: Anatomy of protein pockets and cavities: measurement of binding site geometry and implications for ligand design. Protein science 7(9), 1884–1897 (1998)

31. Lipinski, C.A.: Rule of five in 2015 and beyond: Target and ligand structural limitations, ligand chemistry structure and drug discovery project decisions. Advanced drug delivery reviews 101, 34–41 (2016)

32. Macalino, S.J.Y., Gosu, V., Hong, S., Choi, S.: Role of computer-aided drug design in modern drug discovery. Archives of pharmacal research 38(9), 1686–1701 (2015)

33. Naderi, M., Lemoine, J.M., Govindaraj, R.G., Kana, O.Z., Feinstein, W.P., Brylinski, M.: Binding site matching in rational drug design: Algorithms and applications. Brief Bioinform p. bby078 (2018)

34. Roche, D.B., Brackenridge, D.A., McGuffin, L.J.: Proteins and their interacting partners: An introduction to protein–ligand binding site prediction methods. International journal of molecular sciences 16(12), 29829–29842 (2015)

35. Ronneberger, O., Fischer, P., Brox, T.: U-net: Convolutional networks for biomedical image segmentation. In: International Conference on Medical image computing and computer-assisted intervention. pp. 234–241. Springer (2015)

36. Roy, A., Yang, J., Zhang, Y.: Cofactor: an accurate comparative algorithm for structure-based protein function annotation. Nucleic acids research 40(W1), W471–W477 (2012)

37. Schmidt, T., Haas, J., Cassarino, T.G., Schwede, T.: Assessment of ligand-binding residue predictions in casp9. Proteins: Structure, Function, and Bioinformatics 79(S10), 126–136 (2011)

38. Schmidtke, P., Souaille, C., Estienne, F., Baurin, N., Kroemer, R.T.: Large-scale comparison of four binding site detection algorithms. Journal of chemical information and modeling 50(12), 2191–2200 (2010)

39. Simões, T., Lopes, D., Dias, S., Fernandes, F., Pereira, J., Jorge, J., Bajaj, C., Gomes, A.: Geometric detection algorithms for cavities on protein surfaces in molecular graphics: a survey. In: Computer graphics forum. vol. 36, pp. 643–683. Wiley Online Library (2017)

40. Skolnick, J., Brylinski, M.: Findsite: a combined evolution/structure-based approach to protein function prediction. Briefings in bioinformatics 10(4), 378–391 (2009)

41. Smith, R.D., Clark, J.J., Ahmed, A., Orban, Z.J., Dunbar Jr, J.B., Carlson, H.A.: Updates to binding moad (mother of all databases): Polypharmacology tools and their utility in drug repurposing. Journal of molecular biology 431(13), 2423–2433 (2019)

42. Weisel, M., Proschak, E., Schneider, G.: Pocketpicker: analysis of ligand binding-sites with shape descriptors. Chemistry Central Journal 1(1), 7 (2007)

43. Xu, J., Zhang, Y.: How significant is a protein structure similarity with tm-score= 0.5? Bioinformatics 26(7), 889–895 (2010)

44. Yang, J., Roy, A., Zhang, Y.: Protein–ligand binding site recognition using complementary binding-specific substructure comparison and sequence profile alignment. Bioinformatics 29(20), 2588–2595 (2013)

45. Yu, J., Zhou, Y., Tanaka, I., Yao, M.: Roll: a new algorithm for the detection of protein pockets and cavities with a rolling probe sphere. Bioinformatics 26(1), 46–52 (2009)

46. Yuan, Y., Pei, J., Lai, L.: Binding site detection and druggability prediction of protein targets for structure-based drug design. Current pharmaceutical design 19(12), 2326–2333 (2013)

47. Zhang, Z., Li, Y., Lin, B., Schroeder, M., Huang, B.: Identification of cavities on protein surface using multiple computational approaches for drug binding site prediction. Bioinformatics 27(15), 2083–2088 (2011)

48. Zhou, W., Yan, H.: Alpha shape and delaunay triangulation in studies of protein-related interactions. Briefings in bioinformatics 15(1), 54–64 (2012)

