## Supplemental Materials for PointSite for "PointSite: a point cloud segmentation tool for identification of protein ligand binding atoms"

### 1 LIG_Tool: a tool for extracting biologically relevant ligands from original PDB file:

Similar as P2Rank and DeepSite, PointSite is focused on predicting binding sites for biologically relevant ligands. However, original PDB files in considered datasets often contain more than one such ligand of interest. Moreover, PDB files also contain a variety of other HET groups like modified residues, solvents, salt, large nucleotide molecules, and misplaced groups (which are not in contact with the protein). Although most of the proteins at chain-level contain only one ligand, still there are about 25% contain more than 1 biologically relevant ligands. Therefore, we determine relevant ligands by the following rules:

– number of ligand atoms is greater or equal than 5;
– number of binding residues is greater or equal than 6;
– not consider ligands labeled as 'Poly-Nucleotides';
– filter those ligands that are labeled as 'invalid' according to MOAD database.

For more details about the filter list from MOAD database, please refer to the below link:
`https://github.com/realbigws/LIG_Tool/tree/master/filter_data/moad_filter_data_list`
which contains about 170 ligands.

In summary, simply type the below command to run LIG_Tool for extracting the biologically relevant ligands from a given official PDB file:

```
LIG_Tool -i ${input}.pdb -l 1 -d 6.5 -p ${out_root} -q ${out_root}
-L ${out_root} -O ${out_root} -T 1 -t 'IOPX' -f ${filter_list} -m 6 -N 5
```

where the parameters are explained as follows:
-d: the distance cutoff to define the ligand-binding atoms from the protein, which then forms the ligand binding site (LBS).
-t: select ligand types. Here we select 'IOPX' ligands (I:ions, O:organic, P:peptides, X:others), but no 'N' (Poly-Nucleotides).
-f: the filter list created from MOAD database for those 'invalid' ligands.
-m: mininal number of binding residues.
-N =: minimal number of ligand atoms.

### 2 Detailed description of the testing datasets:

– B277: a dataset containing 277 proteins in a bound state from LIGSITE_csc benchmark released at the year 2006. After LIG_Tool selection, we have 263 valid entries at protein-level in this dataset.
– DT198: a dataset containing 198 drug-target complexes from MetaPocket2 benchmark released at the year 2011. After LIG_Tool selection, we have 163 valid entries at chain-level in this dataset.

- ASTEX85: a dataset containing 85 entries introduced by [astex] at the year 2007, which is used for evaluating molecular docking methods and the ligands all meet with drug-like criteria. After LIG_Tool selection, we have 82 valid entries at protein-level in this dataset.
- CHEN251: a dataset containing 251 entries introduced by [chen11] at the year 2011, which is a non-redundant dataset containing typical representatives at SCOP family level. Note that this dataset has been used to evaluate a variety of LBS identification methods released before the year 2011. After LIG_Tool selection, we have 238 valid entries at chain-level in this dataset.
- COACH420: a dataset containing 420 proteins that contain a mix of drug targets and naturally occurring ligands. This is a subset of the COACH benchmark released at the year 2013, and have no redundancy with B277, DT198, ASTEX85, and CHEN251. After LIG_Tool selection, we have 402 valid entries at chain-level in this dataset.
- HOLO4k: this is a large dataset containing 4543 protein-ligand complexes in the holo form, which has been used to evaluate four popular pocket-centric approaches before the year 2010. Again, it has no redundancy with B277, DT198, ASTEX85, and CHEN251. After LIG_Tool selection, we have 4063 valid entries at protein-level in this dataset.

### 3 More experiment results:

**Table 1.** Comparison of identification performance on B277, DT198, ASTEX85, CHEN251, COACH420 and HOLO4K datasets in terms of atom-IoU criterion with 4.5, 5.5, 6.5Å distance as the identification radius. We also provide the results generated by PointSite.

| Method | *B277* | *DT198* | *ASTEX85* | *CHEN251* | *COACH420* | *HOLO4k* | Metric |
|---|---|---|---|---|---|---|---|
| FPocket | 22.2 | 17.7 | 25.7 | 18.0 | 21.3 | 21.6 | 4.5 |
| | 29.8 | 22.4 | 33.4 | 23.5 | 28.2 | 28.7 | 5.5 |
| | 31.5 | 23.2 | 34.1 | 25.4 | 30.0 | 30.5 | 6.5 |
| | 31.5 | 23.2 | 34.1 | 25.4 | 30.0 | 30.5 | max |
| SiteHound | 26.5 | 18.9 | 27.6 | 21.2 | 24.9 | 24.1 | 4.5 |
| | 34.5 | 23.3 | 36.8 | 27.4 | 32.6 | 32.1 | 5.5 |
| | 36.4 | 23.1 | 38.9 | 29.4 | 34.9 | 34.6 | 6.5 |
| | 36.4 | 23.1 | 38.9 | 29.4 | 34.9 | 34.6 | max |
| MetaPocket2 | 37.3 | 25.8 | 37.5 | 32.8 | 37.7 | 38.8 | 4.5 |
| | 32.6 | 22.2 | 32.4 | 29.9 | 33.4 | 34.4 | 5.5 |
| | 26.6 | 18.1 | 26.1 | 25.2 | 27.4 | 28.3 | 6.5 |
| | 37.3 | 25.8 | 37.5 | 32.8 | 37.7 | 38.8 | max |
| DeepSite | 30.6 | 26.6 | 32.4 | 21.7 | 27.9 | 26.9 | 4.5 |
| | 34 | 29.1 | 37.4 | 26.1 | 33.3 | 32.1 | 5.5 |
| | 33.3 | 27.2 | 37.2 | 27.4 | 33.9 | 33.2 | 6.5 |
| | 34 | 29.1 | 37.4 | 27.4 | 33.9 | 33.2 | max |
| P2Rank | 41.8 | 34.3 | 39.2 | 49.7 | 37.4 | 40.7 | 4.5 |
| | 49.8 | 38.6 | 47.4 | 56.6 | 45.3 | 48.8 | 5.5 |
| | 47.2 | 35.4 | 44.7 | 51.8 | 44.2 | 46.8 | 6.5 |
| | 49.8 | 38.6 | 47.4 | 56.6 | 45.3 | 48.8 | max |
| PointSite | 60.9 | 45.4 | 61.3 | 54.3 | 59.6 | 63.4 | - |

* The numbers represent average value of atom-IoU criterion (i.e., intersection over union of the ground-truth ligand binding atoms and the identified ligand binding

atoms within the TopN ranking list where N is the number of ligands in considered structure).

** As the size of the pseudo surface points (PSP) for different approaches varies, we show the maximal value of atom-IoU generated by the three different identification radii. This maximal value will be regarded as the original value of the certain approach for the following analysis.

**Table 2.** Comparison of identification performance on B277, DT198, ASTEX85, CHEN251, COACH420 and HOLO4K datasets in terms of atom-IoU using FPocket (FP) filtered by PointSite with different parameters (identification radius, ratioPSP). Note that we also put the original value of FP as well as the best results generated by P2Rank.

| Method | *B277* | *DT198* | *ASTEX85* | *CHEN251* | *COACH420* | *HOLO4k* |
|---|---|---|---|---|---|---|
| P2Rank | 49.8 | 38.6 | 47.4 | 56.6 | 45.3 | 48.8 |
| FP (Original) | 31.5 | 23.2 | 34.1 | 25.4 | 30.0 | 30.5 |
| FP (4.5,0.1) | 49.6 | 40.2 | 55.0 | 40.1 | 46.1 | 51.5 |
| FP (4.5,0.2) | 49.6 | 40.1 | 54.9 | 40.2 | 46.1 | 51.5 |
| FP (4.5,0.3) | 49.6 | 39.9 | 54.9 | 40.0 | 46.0 | 51.4 |
| FP (5.5,0.1) | 53.3 | 42.4 | 58.4 | 42.8 | 49.2 | 55.3 |
| FP (5.5,0.2) | 53.2 | 42.1 | 58.4 | 42.7 | 49.1 | 55.2 |
| FP (5.5,0.3) | 53.1 | 42.1 | 58.4 | 42.6 | 49.0 | 55.1 |
| FP (6.5,0.1) | 60.5 | 45.5 | 64.3 | 49.0 | 55.9 | 62.6 |
| FP (6.5,0.2) | 60.5 | 45.1 | 64.3 | 48.8 | 55.8 | 62.6 |
| FP (6.5,0.3) | 60.3 | 45.1 | 64.3 | 48.6 | 55.6 | 62.5 |

**Table 3.** Comparison of identification performance on B277, DT198, ASTEX85, CHEN251, COACH420 and HOLO4K datasets in terms of atom-IoU using FPocket (FP) filtered by PointSite with different parameters (identification radius, ratioPSP). Note that we also put the original value of FP as well as the best results generated by P2Rank.

| Method | *B277* | *DT198* | *ASTEX85* | *CHEN251* | *COACH420* | *HOLO4k* |
|---|---|---|---|---|---|---|
| P2Rank | 49.8 | 38.6 | 47.4 | 56.6 | 45.3 | 48.8 |
| SH (Original) | 36.4 | 23.1 | 38.9 | 29.4 | 34.9 | 34.6 |
| SH (4.5,0.1) | 52.1 | 38.6 | 51.4 | 43.7 | 47.0 | 50.3 |
| SH (4.5,0.2) | 52.3 | 38.7 | 51.4 | 43.7 | 47.0 | 50.3 |
| SH (4.5,0.3) | 52.3 | 38.6 | 51.4 | 43.6 | 47.0 | 50.3 |
| SH (5.5,0.1) | 55.3 | 42.6 | 55.0 | 46.6 | 49.8 | 54.0 |
| SH (5.5,0.2) | 55.3 | 42.6 | 55.0 | 46.5 | 49.8 | 54.0 |
| SH (5.5,0.3) | 55.3 | 42.5 | 55.1 | 46.5 | 49.8 | 54.0 |
| SH (6.5,0.1) | 61.7 | 46.7 | 61.8 | 52.0 | 56.1 | 61.5 |
| SH (6.5,0.2) | 61.7 | 46.6 | 61.8 | 51.9 | 56.1 | 61.5 |
| SH (6.5,0.3) | 61.7 | 46.6 | 61.8 | 51.9 | 55.9 | 61.5 |

**Table 4.** Comparison of identification performance on B277, DT198, ASTEX85, CHEN251, COACH420 and HOLO4K datasets in terms of atom-IoU using FPocket (FP) filtered by PointSite with different parameters (identification radius, ratioPSP). Note that we also put the original value of FP as well as the best results generated by P2Rank.

| Method | *B277* | *DT198* | *ASTEX85* | *CHEN251* | *COACH420* | *HOLO4k* |
|---|---|---|---|---|---|---|
| P2Rank | 49.8 | 38.6 | 47.4 | 56.6 | 45.3 | 48.8 |
| MP (Original) | 37.3 | 25.8 | 37.5 | 32.8 | 37.7 | 38.8 |
| MP (4.5,0.1) | 62.2 | 44.3 | 62.1 | 51.8 | 58.4 | 64.2 |
| MP (4.5,0.2) | 61.9 | 44.3 | 61.3 | 51.5 | 58.3 | 63.9 |
| MP (4.5,0.3) | 61.3 | 43.3 | 61.0 | 50.5 | 57.8 | 62.7 |
| MP (5.5,0.1) | 62.3 | 44.3 | 62.0 | 52.2 | 58.6 | 64.4 |
| MP (5.5,0.2) | 62.1 | 44.2 | 61.3 | 51.8 | 58.1 | 63.9 |
| MP (5.5,0.3) | 60.4 | 42.7 | 60.5 | 51.0 | 57.2 | 61.9 |
| MP (6.5,0.1) | 61.8 | 44.2 | 61.6 | 52.7 | 58.8 | 64.1 |
| MP (6.5,0.2) | 61.5 | 43.8 | 61.1 | 51.9 | 58.2 | 63.2 |
| MP (6.5,0.3) | 58.1 | 41.3 | 58.2 | 49.3 | 54.9 | 58.5 |

**Table 5.** Comparison of identification performance on B277, DT198, ASTEX85, CHEN251, COACH420 and HOLO4K datasets in terms of atom-IoU using DeepSite (DS) filtered by PointSite with different parameters (identification radius, ratioPSP). Note that we also put the original value of DS as well as the best results generated by P2Rank.

| Method | *B277* | *DT198* | *ASTEX85* | *CHEN251* | *COACH420* | *HOLO4k* |
|---|---|---|---|---|---|---|
| P2Rank | 49.8 | 38.6 | 47.4 | 56.6 | 45.3 | 48.8 |
| DS (Original) | 34.0 | 29.1 | 37.4 | 27.4 | 33.9 | 33.2 |
| DS (4.5,0.1) | 43.8 | 35.1 | 42.3 | 30.9 | 38.0 | 36.4 |
| DS (4.5,0.2) | 43.6 | 34.9 | 42.2 | 30.7 | 37.8 | 36.3 |
| DS (4.5,0.3) | 43.5 | 34.9 | 42.1 | 30.3 | 37.7 | 36.1 |
| DS (5.5,0.1) | 47.2 | 37.8 | 45.8 | 33.9 | 41.3 | 39.7 |
| DS (5.5,0.2) | 47.1 | 37.6 | 45.8 | 33.8 | 41.1 | 39.6 |
| DS (5.5,0.3) | 47.0 | 37.4 | 45.7 | 33.3 | 40.9 | 39.4 |
| DS (6.5,0.1) | 52.8 | 39.9 | 51.3 | 39.6 | 47.2 | 45.7 |
| DS (6.5,0.2) | 52.6 | 39.5 | 51.2 | 39.4 | 47.0 | 45.5 |
| DS (6.5,0.3) | 52.4 | 38.9 | 51.1 | 38.6 | 46.6 | 45.1 |

**Table 6.** Comparison of identification performance on B277, DT198, ASTEX85, CHEN251, COACH420 and HOLO4K datasets in terms of atom-IoU using P2Rank (PR) filtered by PointSite with different parameters (identification radius, ratioPSP). Note that we also put the original value of PR.

| Method | *B277* | *DT198* | *ASTEX85* | *CHEN251* | *COACH420* | *HOLO4k* |
|---|---|---|---|---|---|---|
| PR (Original) | 49.8 | 38.6 | 47.4 | 56.6 | 45.3 | 48.8 |
| PR (4.5,0.1) | 56.8 | 43.6 | 56.2 | 54.5 | 50.4 | 57.3 |
| PR (4.5,0.2) | 56.8 | 43.4 | 56.2 | 54.4 | 50.4 | 57.2 |
| PR (4.5,0.3) | 56.8 | 43.4 | 56.3 | 54.4 | 50.4 | 57.1 |
| PR (5.5,0.1) | 60.0 | 45.4 | 59.8 | 56.1 | 53.4 | 60.7 |
| PR (5.5,0.2) | 60.0 | 45.3 | 59.8 | 56.0 | 53.5 | 60.6 |
| PR (5.5,0.3) | 60.0 | 45.3 | 59.8 | 55.9 | 53.4 | 60.5 |
| PR (6.5,0.1) | 63.4 | 46.5 | 62.6 | 57.3 | 57.5 | 64.7 |
| PR (6.5,0.2) | 63.4 | 46.4 | 62.6 | 57.2 | 57.5 | 64.7 |
| PR (6.5,0.3) | 63.5 | 46.0 | 61.2 | 56.8 | 57.3 | 64.3 |

**Table 7.** Comparison of identification performance on B277, DT198, ASTEX85, CHEN251, COACH420 and HOLO4K datasets in terms of DCA using FPocket (FP) filtered by PointSite with different parameters (identification radius, ratioPSP). Note that we also put the original value of FP as well as the best results generated by P2Rank.

| Method | *B277* | *DT198* | *ASTEX85* | *CHEN251* | *COACH420* | *HOLO4k* |
|---|---|---|---|---|---|---|
| P2Rank | 74.9 | 68.6 | 65.1 | 79.2 | 68.9 | 75.1 |
| FP (Original) | 44.0 | 32.8 | 43.4 | 34.2 | 43.6 | 42.4 |
| FP (4.5,0.1) | 75.3 | 63.8 | 73.6 | 58.7 | 72.9 | 79.3 |
| FP (4.5,0.2) | 75.0 | 63.3 | 73.6 | 58.7 | 72.7 | 79.3 |
| FP (4.5,0.3) | 75.3 | 63.3 | 73.6 | 58.0 | 72.5 | 79.2 |
| FP (5.5,0.1) | 74.5 | 64.4 | 73.6 | 59.1 | 72.7 | 80.0 |
| FP (5.5,0.2) | 74.2 | 63.8 | 73.6 | 58.9 | 72.5 | 79.9 |
| FP (5.5,0.3) | 74.5 | 63.8 | 73.6 | 58.4 | 72.1 | 79.8 |
| FP (6.5,0.1) | 75.0 | 63.8 | 73.6 | 59.1 | 73.0 | 80.2 |
| FP (6.5,0.2) | 74.5 | 63.3 | 73.6 | 58.9 | 72.9 | 80.1 |
| FP (6.5,0.3) | 74.2 | 63.3 | 73.6 | 58.2 | 71.8 | 79.9 |

**Table 8.** Comparison of identification performance on B277, DT198, ASTEX85, CHEN251, COACH420 and HOLO4K datasets in terms of DCA using SiteHound (SH) filtered by PointSite with different parameters (identification radius, ratioPSP). Note that we also put the original value of SH as well as the best results generated by P2Rank.

| Method | *B277* | *DT198* | *ASTEX85* | *CHEN251* | *COACH420* | *HOLO4k* |
|---|---|---|---|---|---|---|
| P2Rank | 74.9 | 68.6 | 65.1 | 79.2 | 68.9 | 75.1 |
| SH (Original) | 53.9 | 39.0 | 54.3 | 45.6 | 53.2 | 53.1 |
| SH (4.5,0.1) | 78.1 | 65.5 | 76.0 | 67.9 | 76.8 | 85.6 |
| SH (4.5,0.2) | 78.4 | 65.5 | 76.0 | 67.4 | 76.8 | 85.5 |
| SH (4.5,0.3) | 78.6 | 65.5 | 76.0 | 67.2 | 76.4 | 85.5 |
| SH (5.5,0.1) | 78.6 | 70.1 | 76.0 | 68.5 | 76.4 | 86.3 |
| SH (5.5,0.2) | 78.6 | 70.1 | 76.0 | 68.3 | 76.4 | 86.3 |
| SH (5.5,0.3) | 78.6 | 70.1 | 76.0 | 68.3 | 76.4 | 86.2 |
| SH (6.5,0.1) | 78.4 | 70.6 | 76.7 | 67.4 | 76.8 | 86.6 |
| SH (6.5,0.2) | 78.4 | 70.6 | 76.7 | 67.2 | 77.0 | 86.6 |
| SH (6.5,0.3) | 78.4 | 70.6 | 76.7 | 67.2 | 76.3 | 86.4 |

**Table 9.** Comparison of identification performance on B277, DT198, ASTEX85, CHEN251, COACH420 and HOLO4K datasets in terms of DCA using MetaPocket2 (MP) filtered by PointSite with different parameters (identification radius, ratioPSP). Note that we also put the original value of MP as well as the best results generated by P2Rank.

| Method | *B277* | *DT198* | *ASTEX85* | *CHEN251* | *COACH420* | *HOLO4k* |
|---|---|---|---|---|---|---|
| P2Rank | 74.9 | 68.6 | 65.1 | 79.2 | 68.9 | 75.1 |
| MP (Original) | 60.9 | 49.1 | 59.7 | 46.8 | 59.2 | 60.1 |
| MP (4.5,0.1) | 74.2 | 62.6 | 67.2 | 56.8 | 68.2 | 70.9 |
| MP (4.5,0.2) | 73.4 | 62.0 | 68.1 | 56.6 | 68.2 | 71.3 |
| MP (4.5,0.3) | 73.4 | 62.6 | 68.9 | 55.9 | 67.9 | 71.0 |
| MP (5.5,0.1) | 74.2 | 62.6 | 67.2 | 57.5 | 68.4 | 71.3 |
| MP (5.5,0.2) | 73.7 | 62.0 | 68.1 | 56.6 | 68.4 | 71.5 |
| MP (5.5,0.3) | 72.7 | 61.3 | 68.9 | 56.1 | 67.7 | 70.7 |
| MP (6.5,0.1) | 74.0 | 62.6 | 67.2 | 57.7 | 69.1 | 71.6 |
| MP (6.5,0.2) | 73.7 | 62.6 | 68.1 | 56.1 | 68.4 | 71.4 |
| MP (6.5,0.3) | 71.6 | 62.0 | 63.9 | 54.3 | 66.1 | 68.2 |

**Table 10.** Comparison of identification performance on B277, DT198, ASTEX85, CHEN251, COACH420 and HOLO4K datasets in terms of DCA using DeepSite (DS) filtered by PointSite with different parameters (identification radius, ratioPSP). Note that we also put the original value of DS as well as the best results generated by P2Rank.

| Method | *B277* | *DT198* | *ASTEX85* | *CHEN251* | *COACH420* | *HOLO4k* |
|---|---|---|---|---|---|---|
| P2Rank | 74.9 | 68.6 | 65.1 | 79.2 | 68.9 | 75.1 |
| DS (Original) | 59.9 | 52.6 | 60.2 | 43.4 | 55.0 | 58.8 |
| DS (4.5,0.1) | 66.1 | 59.4 | 61.7 | 47.7 | 59.5 | 62.3 |
| DS (4.5,0.2) | 65.6 | 59.4 | 61.7 | 47.7 | 59.7 | 62.2 |
| DS (4.5,0.3) | 65.6 | 59.4 | 61.7 | 47.3 | 59.5 | 62.0 |
| DS (5.5,0.1) | 66.7 | 61.1 | 61.7 | 47.7 | 59.5 | 62.4 |
| DS (5.5,0.2) | 66.4 | 61.1 | 61.7 | 47.7 | 59.7 | 62.2 |
| DS (5.5,0.3) | 66.4 | 61.1 | 61.7 | 47.0 | 59.5 | 62.0 |
| DS (6.5,0.1) | 66.9 | 60.6 | 61.7 | 47.9 | 59.7 | 62.5 |
| DS (6.5,0.2) | 66.7 | 60.6 | 61.7 | 47.9 | 59.5 | 62.3 |
| DS (6.5,0.3) | 66.7 | 60.6 | 61.7 | 47.3 | 59.5 | 62 .0 |

**Table 11.** Comparison of identification performance on B277, DT198, ASTEX85, CHEN251, COACH420 and HOLO4K datasets in terms of DCA using P2Rank (PR) filtered by PointSite with different parameters (identification radius, ratioPSP). Note that we also put the original value of PR.

| Method | *B277* | *DT198* | *ASTEX85* | *CHEN251* | *COACH420* | *HOLO4k* |
|---|---|---|---|---|---|---|
| PR (Original) | 74.9 | 68.6 | 65.1 | 79.2 | 68.9 | 75.1 |
| PR (4.5,0.1) | 77.5 | 72.0 | 70.5 | 71.8 | 71.2 | 77.3 |
| PR (4.5,0.2) | 77.3 | 72.0 | 70.5 | 71.1 | 71.0 | 80.8 |
| PR (4.5,0.3) | 77.3 | 72.6 | 70.5 | 70.0 | 70.9 | 80.6 |
| PR (5.5,0.1) | 77.5 | 73.7 | 70.5 | 71.6 | 71.2 | 81.0 |
| PR (5.5,0.2) | 77.5 | 73.7 | 70.5 | 70.7 | 71.0 | 80.9 |
| PR (5.5,0.3) | 77.5 | 73.7 | 70.5 | 70.0 | 70.9 | 80.6 |
| PR (6.5,0.1) | 77.8 | 74.9 | 70.5 | 71.3 | 71.2 | 81.0 |
| PR (6.5,0.2) | 77.5 | 74.9 | 70.5 | 70.2 | 71.0 | 80.8 |
| PR (6.5,0.3) | 77.5 | 74.9 | 69.8 | 68.8 | 70.9 | 80.3 |
